# Adenosine receptor and its downstream targets, mod(mdg4) and Hsp70, work as a signaling pathway modulating cytotoxic damage in *Drosophila*

**DOI:** 10.1101/2019.12.22.886143

**Authors:** Yu-Hsien Lin, Houda Ouns Maaroufi, Lucie Kucerova, Lenka Rouhova, Tomas Filip, Michal Zurovec

**Author notes:** Correspondence and requests for materials should be addressed to Y.H.L. and M.Z. Department of Plant Physiology, Swammerdam Institute for Life Sciences, University of Amsterdam, 1098 XH Amsterdam, The Netherlands.

## Abstract

Adenosine (Ado) is an important signaling molecule involved in stress responses. Studies in mammalian models have shown that Ado regulates signaling mechanisms involved in ‘danger-sensing’ and tissue-protection. Yet, little is known about the role of Ado signaling in *Drosophila*. In the present study, we observed lower extracellular Ado concentration and suppressed expression of Ado transporters in flies expressing mutant huntingtin protein (mHTT). We altered Ado signaling using genetic tools and found that the overexpression of Ado metabolic enzymes, as well as the suppression of Ado receptor (AdoR) and transporters (ENTs), were able to minimize mHTT-induced mortality. We also identified the downstream targets of the AdoR pathway, the modifier of mdg4 (Mod(mdg4)) and heat-shock protein 70 (Hsp70), which carry out its function. Finally, we showed that a decrease in Ado signaling affect other *Drosophila* stress reactions, including paraquat and heat-shock treatments. Our study provides important insights into how Ado regulates stress responses in *Drosophila*.

## Introduction

Tissue injury, ischemia, and inflammation activate organismal responses involved in the maintenance of tissue homeostasis. Such responses require precise coordination among the involved signaling pathways. Adenosine (Ado) represents one of the key signals contributing to the orchestration of cytoprotection, immune reactions, and regeneration, as well as balancing energy metabolism (Borea et al., 2016). Under normal conditions, the Ado concentration in blood is in the nanomolar range; however, under pathological circumstances the extracellular Ado (e-Ado) level may dramatically change (Moser et al., 1989). Ado has previously been considered a retaliatory metabolite, having general tissue protective effects. Prolonged adenosine signaling, however, can exacerbate tissue dysfunction in chronic diseases (Antonioli et al., 2019) Ado therefore functions as an imperfect protector; in some cases it may be beneficial, while in others it may worsen tissue damage (Picano and Abbracchio, 2000).

Ado signaling is well-conserved among phyla. The concentration of Ado in the *D. melanogaster* hemolymph is maintained in the nanomolar range, as in mammals, and increases dramatically in adenosine deaminase mutants or during infections (Dolezelova et al., 2005;Novakova and Dolezal, 2011). Unlike mammals, *D. melanogaster* contains only a single AdoR isoform (stimulating cAMP) and several proteins that have Ado metabolic and transport activities involved in the fine regulation of adenosine levels. *Drosphila melanogaster* adenosine deaminase-related growth factors (ADGFs), which are related to human ADA2, together with adenosine kinase (AdenoK) are the major metabolic enzymes converting extra- and intra-cellular adenosine to inosine and AMP, respectively (Zurovec et al., 2002;Maier et al., 2005;Stenesen et al., 2013). The transport of Ado across the plasma membrane is mediated by three equilibrative and two concentrative nucleoside transporters (ENTs and CNTs, respectively) similar to their mammalian counterparts. Ado signaling in *Drosophila* has been reported to affect various physiological processes, including the regulation of synaptic plasticity in the brain, proliferation of gut stem cells, hemocyte differentiation, and metabolic adjustments during the immune response (Knight et al., 2010;Mondal et al., 2011;Bajgar et al., 2015;Xu et al., 2020).

The present study examined the role of *Drosophila* Ado signaling on cytotoxic stress and aimed to elicit the underlying mechanism. Earlier reports have shown that expression of the expanded polyglutamine domain from human mutant huntingtin protein (mHTT) induces cell death in both *Drosophila* neurons and hemocytes (Marsh et al., 2000;Lin et al., 2019). In our study, we confirmed the low-viability phenotype of mHTT-expressing larvae and observed that such larvae display a lower level of e-Ado in the hemolymph. Furthermore, we used genetic tools and altered the expression of genes involved in Ado metabolism and transport to find out whether changes in Ado signaling can modify the phenotype of mHTT-expressing flies. Finally, we uncovered a downstream mechanism of the *Drosophila* Ado pathway, namely *mod(mdg4)* and Hsp70, which modify both the formation of mHTT aggregates and the stress response to heat-shock and paraquat treatments.

## Results

### Decreased hemolymph Ado titer in mHTT-expressing larvae

To characterize the involvement of Ado signaling in the stress response, we used mHTT-expressing flies as a well-characterized genetic model for neurodegeneration and cytotoxic stress (Rosas-Arellano et al., 2018). We initially examined flies overexpressing normal exon 1 from human huntingtin (Q20 HTT), or its mutant pathogenic form (Q93 mHTT), driven by the ubiquitous *daughterless-Gal4* (*da-Gal4*) and pan-neuron driver (*elav-Gal4*). We observed that 100% of Q93-expressing larvae driven by *da-Gal4* died during the wandering stage. In contrast, those driven by *elav-Gal4* displayed no impact on larval development (Fig. S1A) but with a reduced adult eclosion rate (Fig. S1B) and lifespan (Fig. S1C). These results are consistent with previous observations (Song et al., 2013).

Measurement of the extracellular Ado (e-Ado) concentration in the hemolymph of Q93-expressing larvae (3^rd^ instar) showed that its level was significantly lower compared to larvae expressing Q20 or control *da-GAL4* only (Fig. 1A). Since e-Ado concentration may be associated with the level of extracellular ATP (e-ATP), we also examined its titer in larval hemolymph. However, as shown in Fig. 1B, there was no significant difference in e-ATP levels between Q20, Q93, and control *da-GAL4* larvae.

**Figure 1.**
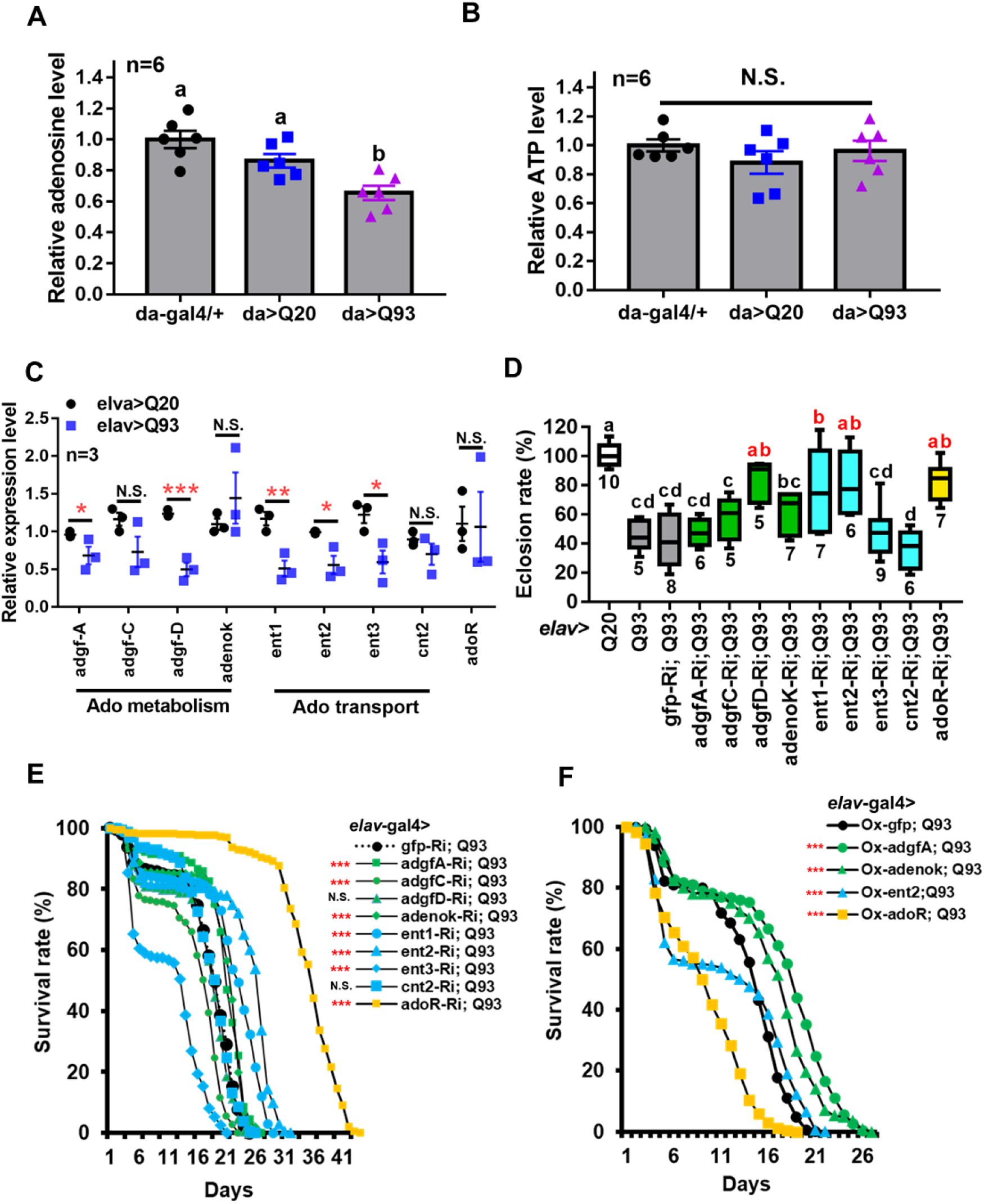
Reduced extracellular Ado transport and receptor suppress mHTT induced lethality. (A-B) Relative level of extracellular Ado (A) and ATP (B) titers in Q93-expressing (da>Q93), Q20-expressing (da>Q20), and control *da*-GAL4 (da/+) larvae. Ado and ATP concentration are normalized to control larvae. Significance was analyzed by ANOVA; significant differences (*P* < 0.05) among treatment groups are marked with different letters; N.S., not significant; n = 6. Error bars are presented as mean ± SEM (C) Transcriptions level of genes involved in regulating Ado homeostasis in Q93-expressing (elav>Q93) and control Q20-expressing (elav>Q20) larval brains. Significance was analyzed by Student’s t-test and labeled as follows: \**P* < 0.05, \*\**P* < 0.01, \*\*\**P* < 0.001; N.S., not significant. n = 3. Error bars are presented as mean ± SEM (D) Eclosion rate of mHTT-expressing adult females (elav>Q93) with RNAi silencing (Ri) Ado metabolic enzymes, transporters, *adoR* and control *gfp*. Numbers below each column indicate the number of replicates (n). Significance was analyzed by ANOVA; significant differences (*P* < 0.05) among treatment groups are marked with different letters (E) Survival of mHTT-expressing adult females (elav>Q93) with RNAi silencing (Ri) Ado metabolic enzymes, transporters, *adoR* and control *gfp*. Significance was analyzed by weighted log-rank test; significant differences between each treatment group and control (*gfp*-Ri) are labeled as follows: \*\*\**P* < 0.001; N.S., not significant. n > 200 (F) Survival of mHTT-expressing adult females (elav>Q93) overexpressing (Ox) Ado metabolic enzymes (*adgf-A* and *adenok*), transporters (*ent2*), *adoR*, and control *gfp*. Significance was analyzed by a weighted log-rank test; significant differences between each treatment group to control (*gfp*-Ri) are labeled as \*\*\**P* < 0.001. n > 200

We thus postulated that the lower level of e-Ado in Q93 larvae might be caused by changes in genes involved in Ado metabolism or transportation. Therefore, we compared the expression of *adgf* genes (*adgf-a*, *adgf-c*, *adgf-d*), adenosine kinase (*adenoK*), adenosine transporters (*ent1*, *ent2*, *ent3*, *cnt2*), and adenosine receptor (*adoR*) in the brains of Q93- and Q20-expressing larvae driven by *elav*-*Gal4* (Fig. 1C). The results showed that the expression levels of *adgf-a* and *adgf-d*, as well as transporters *ent1*, *ent2*, and *ent3*, in the brain of Q93 larvae were significantly lower than in Q20 larvae. There was no difference in the expression of *cnt2* and *adoR* between Q93 and Q20 larvae.

### Enhanced e-Ado signaling increased mortality of mHTT flies

To study the effect of e-Ado signaling on mHTT-induced cytotoxicity, we compared the survival of transgenic lines that co-express RNAi constructs of Ado metabolic, transport and receptor genes together with Q93 and Q20 driven by *elav*-*GAL4*. The results showed that knocking down *adgf-D*, *ent1*, *ent2*, and *adoR* resulted in a significantly increased eclosion rate (Fig. 1D), and silencing *adgf-A* and *adenoK*, *ent1*, *ent2*, and *adoR* significantly extended the adult lifespan of mHTT-expressing flies (Fig. 1E). Notably, the RNAi silencing of *ent2* and *adoR* extended the lifespan of mHTT-expressing flies to 30 and 40 days, respectively, which is about 1.5~2 times longer than that of control *gfp*-RNAi-expressing mHTT flies. To ensure that the mortality of the Q93 flies was mainly caused by mHTT expression and not by the RNAi constructs, we examined the survival of flies co-expressing normal htt Q20 together with RNAi transgenes until all corresponding experimental flies (expressing Q93 together with RNAi constructs) died. We did not observe a significant effect for any of the RNAi transgenes on adult survival (Fig. S2).

It is generally assumed that gain- and loss-of-function manipulations of functionally important genes should lead to the opposite phenotypes. We therefore tested whether the overexpression of *adgf-A*, *adenoK*, *ent2*, and *adoR* would rescue mHTT phenotypes. As shown in Fig. 1F, increasing either the intra- or extracellular Ado metabolism by overexpressing *adenoK* and *adgf-A* in Q93 flies extended their lifespan in comparison to control Q93 flies overexpressing GFP protein. In contrast, the overexpression of *ent2* and *adoR* significantly decreased the lifespan of mHTT-expressing flies. Therefore, the overexpression of *adoR* and *ent2* genes resulted in a phenotype opposite to that observed in the knockdowns, thus supporting the importance of these genes as key regulators of mHTT phenotypes.

### Knocking down *ent2* and *adoR* reduced cell death and mHTT aggregate formation

To determine whether the reduction of Ado signaling could affect other phenotypes of Q93 flies, we examined the effect of knocking down genes involved in Ado signaling and metabolism on *Drosophila* rhabdomere degeneration and mHTT aggregate formation. We expressed RNAi transgenes in the eyes of Q93 flies using the *gmr*-*GAL4* driver (Mugat et al., 2008;Kuo et al., 2013) and compared the levels of retinal pigment cell degeneration (Fig. 2A). The results revealed that silencing Ado metabolic enzymes did not significantly influence the level of retinal pigment cell degeneration; however, retinal pigment cell degeneration was significantly reduced in *ent2* knockdown flies. Surprisingly we did not observe a significant rescue of cell death by silencing *adoR* (Fig S3). We therefore assumed that it might be due to insufficient RNAi efficiency for suppressing AdoR signaling in the eye. To test this, we examined two combinations: mHTT-expressing flies with the adoR RNAi transgene under an adoR heterozygous mutant background (AdoR^1^/+), and mHTT-expressing flies under an AdoR^1^ homozygous mutant background. As shown in Fig. 2A, both had significantly rescued retinal pigment cell degeneration, similar to that of *ent2* RNAi flies.

**Figure 2.**
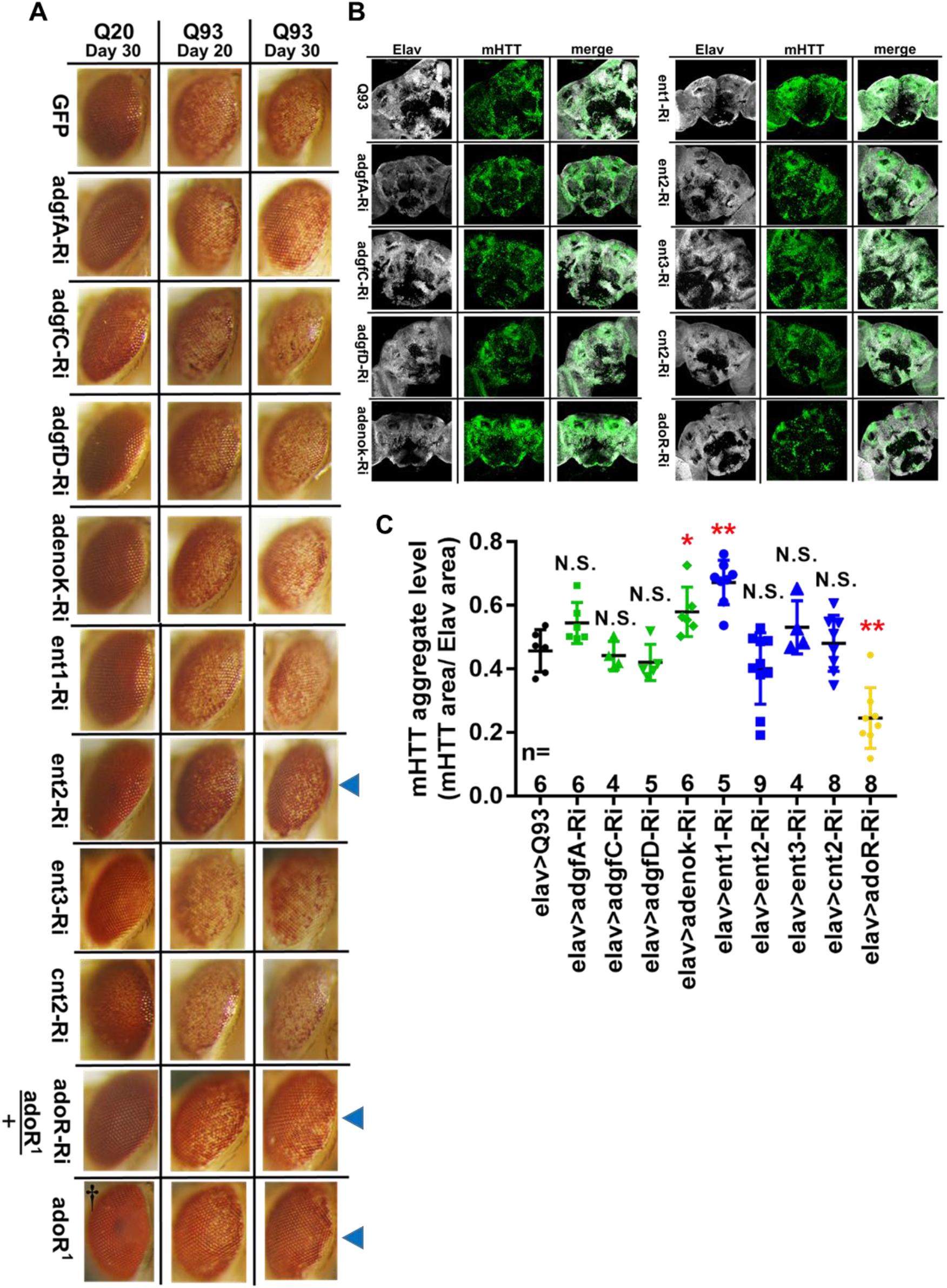
Suppression of *ent2* and *adoR* decreased mHTT-induced cytotoxicity and mHTT aggregate formation. (A) Retinal pigment cell degeneration in mHTT-expressing adult females (gmr>Q93) with RNAi silencing Ado metabolic enzymes, transporters, *adoR* (*adoR* heterozygous mutant background) and mHTT-expressing flies under *adoR* homozygous mutant background. Blue arrows indicate treated groups showing a significantly reduced loss of pigment. †Eye image of control homozygous *adoR^1^* mutant without *htt* expression. Detailed methodologies for sample collection and eye imaging are described in Materials and methods (B) Representative confocal images of the brains of 10-day-old mHTT-expressing adult females (elav>Q93) with RNAi silencing Ado metabolic enzymes, transporters, and *adoR*. Neuronal cells were detected with anti-Elav; mHTT aggregates were detected with anti-HTT (MW8) (C) Level of mHTT aggregation formation was calculated by normalizing the area of mHTT signal to the area of Elav signal. Significance in mHTT aggregate levels was analyzed using a Mann-Whitney U-test; significant differences between control Q93 flies and each RNAi treatment group are labeled as follows: \**P* < 0.05; \*\**P* < 0.01; N.S., not significant. Error bars are presented as mean ± SEM. The number (n) of examined brain images are shown below each bar

To examine the level of mHTT aggregate formation in the *Drosophila* brain, we drove the expression of transgenes using *elav-GAL4* and stained the brains with mHTT antibody (MW8), which exclusively stains mHTT inclusions (Ko et al., 2001). The results showed that mHTT inclusions were reduced to 50% in 10-day-old Q93 *adoR* RNAi flies (Fig. 2B&C), with 20-day-old Q93 *adoR* RNAi flies exhibiting a similar level of suppression (Fig. S4). Our results demonstrate that decreased e-Ado signaling by either knocking down the transporter *ent2* or *adoR* has a strong influence on reducing mHTT-induced cell cytotoxicity and mHTT aggregate formation.

### Epistatic interaction of *adoR* and *ent2* on mHTT-induced mortality

The above results indicated that knockdown of *adoR*, *ent1*, or *ent2* expression significantly extended the adult longevity of mHTT files (Fig. 1E). Therefore, we next tested whether there is a synergy between the effects of *adoR* and both transporters. First, we co-expressed *adoR* RNAi constructs with *ent1* RNAi in Q93-expressing flies. As shown in Fig. 3A, the double knockdown of *ent1* and *adoR* shows a sum of individual effects on lifespan which is greater than the knockdown of *adoR* alone. There seems to be a synergy between *ent1* and *adoR*, suggesting that *ent1* may have its own effect which is partially independent from *adoR* signaling. In contrast, when we performed a double knockdown of *adoR* and *ent2* RNAi in Q93-expressing flies, the silencing of both had the same effect as silencing *adoR* only, indicating that they are involved in the same pathway.

**Figure 3.**
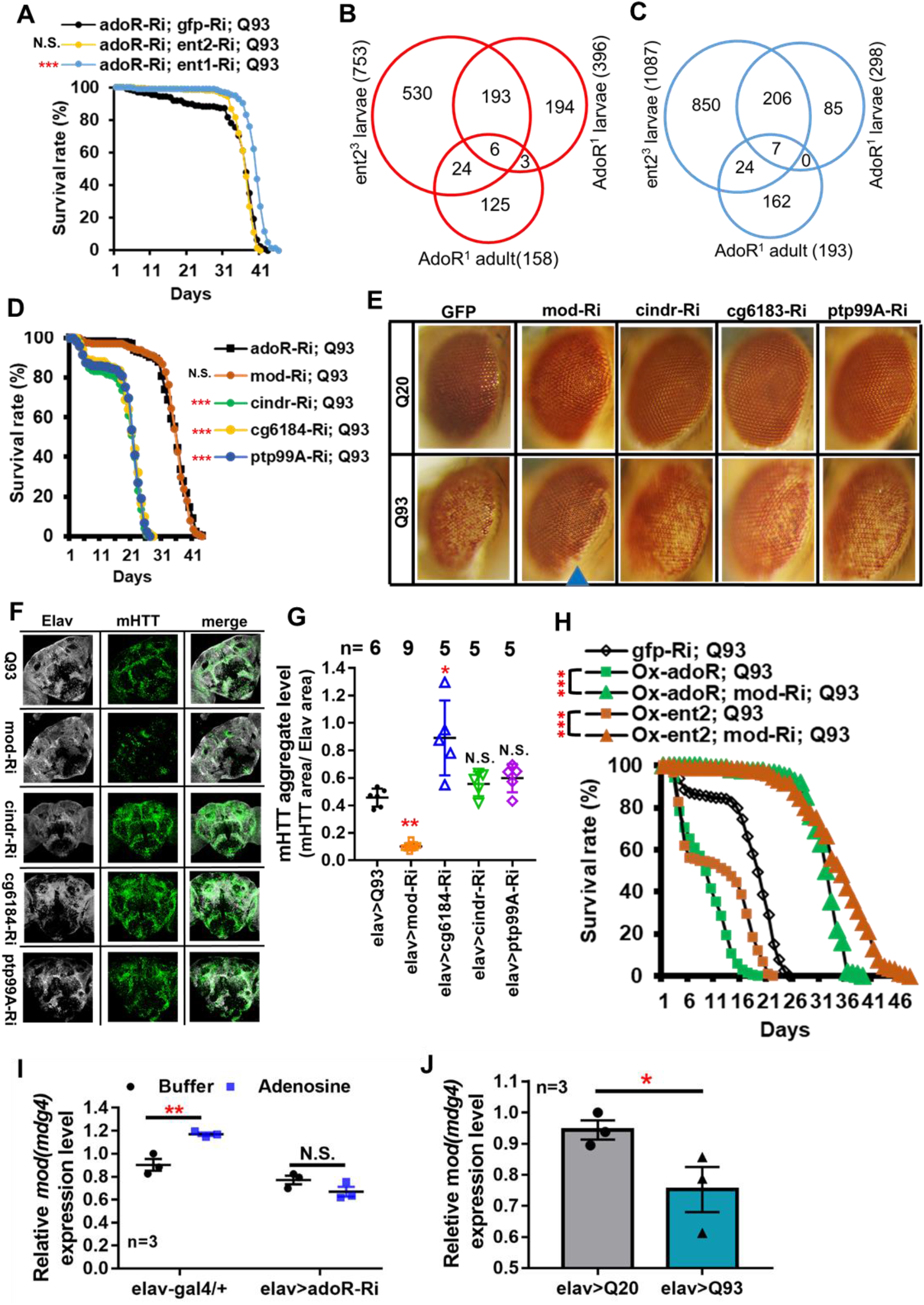
Mod(mdg4) as a downstream target of ENT2/AdoR pathway modulated mHTT effects and aggregate formation. (A) Survival of mHTT-expressing adult females (elav>Q93) with RNAi co-silencing (Ri) transporters (*ent1* or *ent2*) and *adoR*. Significance was analyzed by a weighted log-rank test; significant differences between each treatment group and control (*gfp*-Ri) are labeled as follows: \*\*\**P* < 0.001; N.S., not significant. n > 200 (B-C) Microarray analysis of the transcriptomes of *ent2* and *adoR* mutants. Venn diagram shows the number of common genes (in intersect region) which were upregulated (B) or downregulated (C) among the *adoR* mutant larvae vs. control (w^1118^), *adoR* mutant adults vs. control (w^1118^), and ent2 mutant larvae vs. control (w^1118^). The cutoff values for expression differences were set at Q < 0.05 (false discovery rate, FDR) (D) Survival of mHTT-expressing adult females (elav>Q93) with RNAi co-silencing (Ri) of potential downstream genes of the ENT2/AdoR pathway. Significance was analyzed by a weighted log-rank test; significant differences between each treatment group to control (*adoR*-Ri) are labeled as follows: \*\*\**P* < 0.001; N.S., not significant. n > 200 (E) Retinal pigment cell degeneration in mHTT-expressing adult females (gmr>Q93) with RNAi silencing potential downstream genes of the ENT2/AdoR pathway. Blue arrows indicate treated groups showing a significantly reduced loss of pigment. Detailed methodologies for sample collection and eye imaging are described in Materials and methods (F) Representative confocal images of the brains of 10-day-old mHTT-expressing adult females (elav>Q93) with RNAi silencing potential downstream genes of the ENT2/AdoR pathway. Neuronal cells were detected with anti-Elav and mHTT aggregates were detected with anti-HTT (MW8) (G) The level of mHTT aggregation formation was calculated by normalizing the area of mHTT signal to the area of Elav signal. Significance in mHTT aggregate levels was analyzed using a Mann-Whitney U-test; significant differences between control Q93 flies and each RNAi treatment group are labeled as follows: \**P* < 0.05; \*\**P* < 0.01; N.S., not significant. Error bars are presented as mean ± SEM. The number (n) of examined brain images are indicated above each bar (H) Survival of mHTT-expressing adult females (elav>Q93) with co-RNAi silencing *mod(mdg4)* and co-overexpressing *adoR* or *ent2*. Significance was analyzed by a weighted log-rank test; significant differences are labeled as \*\*\**P* < 0.001. n > 200 (I) Transcription level of *mod(mdg4)* two hours after Ado injection into the whole body of three- to five-day old control adult females (elav-gal4/+) and *adoR* RNAi females (elav>adoR-Ri). Significance was analyzed by Student’s t-test and labeled as follows: \*\**P* < 0.01; N.S., not significant. n = 3. Error bars are presented as mean ± SEM (J) Transcriptions level of *mod(mdg4)* in Q93-expressing (elav>Q93) and control Q20-expressing (elav>Q20) larval brains. Significance was analyzed by Student’s t-test and labeled as \**P* < 0.05. n = 3. Error bars are presented as mean ± SEM

### Identification of potential downstream targets of the AdoR pathway

Our results indicate that *ent2* and *adoR* modify mHTT cytotoxicity and belong to the same pathway. To identify their potential downstream target genes, we compared the gene expression profiles of larvae carrying mutations in *adoR* or *ent2* as well as adult *adoR* mutants by using microarrays (Affymetrix). The data are presented as Venn diagrams, which show the intersection between differentially-expressed genes for individual mutants in all three data sets, including six upregulated (Fig. 3B) and seven downregulated mRNAs (Fig. 3C). According to Flybase annotations (http://flybase.org), four of these genes were expressed in the nervous system (*ptp99A* was upregulated, while *CG6184*, *cindr*, and *mod(mdg4)* were downregulated) (Tab. S1).

In order to examine the potential roles of these four genes in the interaction with mHTT, we co-expressed RNAi constructs of these candidate genes with mHTT and assessed the adult lifespan (Fig. 3D). The results showed that only the knockdown of *mod(mdg4)* extended the lifespan of mHTT-expressing flies, and that the survival curve was not significantly different from that of *adoR* RNAi Q93 flies. Furthermore, *mod(mdg4)* RNAi was the only one of these constructs that significantly reduced retinal pigment cell degeneration (Fig. 3E) and decreased the formation of mHTT inclusions (Fig. 3F&G).

We next examined the possible epistatic relationship between *ent2*, *adoR*, and *mod(mdg4)* by combining the overexpression of *ent2* or *adoR* with *mod(mdg4)* RNAi in mHTT-expressing flies (Fig. 3H). The results showed that the knockdown of *mod(mdg4)* RNAi was able to minimize the lethal effects caused by *ent2* and *adoR* overexpression in mHTT flies. This indicated that *mod(mdg4)* is a downstream target of the AdoR pathway. In addition, we found that increasing the e-Ado concentration by microinjecting Ado significantly increased *mod(mdg4)* expression in GAL4 control flies but not in the flies with *adoR* knockdown (Fig. 3I). *mod(mdg4)* expression in the brain of mHTT Q93 larvae was lower than in control Q20 HTT larvae (Fig. 3J). This result is consistent with a lower e-Ado level in Q93 mHTT larvae (Fig. 1A).

Taken together, our results demonstrate that *mod(mdg4)* serves as a major downstream target of the AdoR pathway, modulating the process of mHTT inclusion formation and mHTT-induced cytotoxicity.

### AdoR pathway with Mod(mdg4) as regulators of Hsp70 protein production

Earlier studies on *Drosophila* protein two-hybrid screening have indicated that Mod(mdg4) is able to interact with six proteins from the Hsp70 family (Giot et al., 2003;Oughtred et al., 2019). In addition, Hsp70 family proteins are known to contribute to suppressing mHTT aggregate formation (Warrick et al., 1999;Chan et al., 2000). In the present study, we compared the levels of Hsp70 protein in *adoR* and *mod(mdg4)* RNAi flies (Figs. 4A, 4B and S5); the results showed that both knockdowns doubled the level of Hsp70 compared to *elav-Gal4* control flies under a non-stress condition (i.e. without mHTT expression). We next compared the level of Hsp70 in flies co-expressing mHTT with each RNAi construct (Figs 4C, 4D and S6). Interestingly, both *adoR* and *mod(mdg4)* RNAi flies co-expressing Q93 mHTT again showed levels around two-fold higher than the Q20 HTT-expressing control, although it was around ten times higher in Q93 mHTT-only flies. These results indicate that *adoR* and *mod(mdg4)* are able to suppress Hsp70 protein production under a non-stress condition. The knockdown of *adoR* and *mod(mdg4)* leads to an increase of Hsp70 production, thus preventing mHTT aggregate formation and decreasing mHTT cytotoxicity.

**Figure 4.**
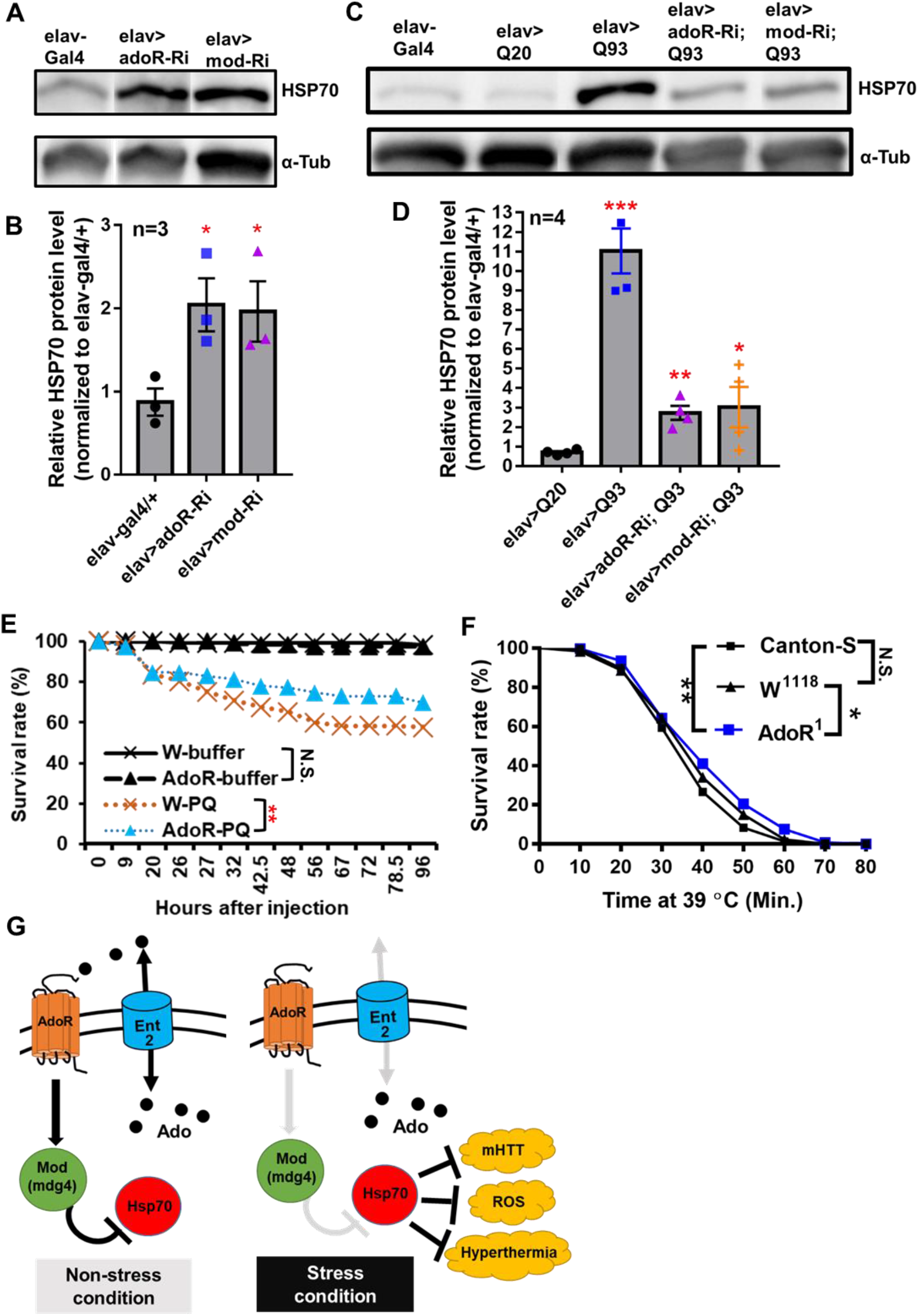
AdoR regulated the Hsp70 protein level and influenced the stress response to paraquat and heat-shock treatments. (A-B) Representative images of western blot analysis. (A) Hsp70 protein level in the head of 10-day-old adult females with RNAi silencing *adoR* (elav>adoR-Ri), *mod(mdg4)* (elav>mod-Ri) and control (elav-gal4/+). (B) The Hsp70 protein level was quantified by normalizing the intensity of the Hsp70 band to the α-Tubulin band using ImageJ; values of RNAi treatment groups were further normalized to the elav-gal4 control. Significance was analyzed by Student’s t-test; significant differences between the control and each RNAi treatment group are labeled as \**P* < 0.05. n = 3. Error bars are presented as mean ± SEM. Original gel images are presented in the supplementary materials (Fig. S6) (C-D) Representative image of western blot analysis. (C) Hsp70 protein level in the head of 10-day-old HTT (elav>Q20) or mHTT expressing (elav>Q93) adult females with RNAi silencing *adoR* and *mod(mdg4)*. (D) The Hsp70 protein level was quantified by normalizing the intensity of the Hsp70 band to the α-Tubulin band by using ImageJ; values of each treatment group were further normalized to the elav-gal4 control. Significance was analyzed by Student’s t-test; significant differences between HTT-expressing flies (elav>Q20) and each RNAi treatment of Q93-expressing flies are labeled as follows: \**P* < 0.05, \*\**P* < 0.01, \*\*\**P* < 0.001. n = 4. Error bars are presented as mean ± SEM. Original gel images are presented in the supplementary materials (Fig. S7) (E) Survival of w^1118^ and homozygous *adoR* mutant adult males after paraquat (PQ) injection. Control groups were injected with ringer buffer. Significance was analyzed by weighted log-rank test; significant differences are labeled as follows: \*\**P* < 0.01, N.S., not significant. W-ringer, n =116; AdoR^1^-ringer, n = 118; W-PQ, n =118; AdoR^1^-PQ, n = 119. (F) Survival of Cantons-S, w^1118^ and homozygous *adoR* mutant (AdoR^1^) adult males during heat-shock treatment. Significance was analyzed by weighted log-rank test; significant differences are labeled as follows: \**P* < 0.05, \*\**P* < 0.01. Cantons-S and W^1118^, n = 300; AdoR^1^, n = 370 (G) Summary model of Ado signaling under stress response. Under a non-stress condition, the activated AdoR and Mod(mdg4) reduce Hsp70 production. In contrast, decreased Ado signaling under a stress condition resulted in Hsp70 production, which in turn enhanced stress tolerance

### Decreased susceptibility to oxidative and heat-shock stresses in *adoR* mutant flies

Since Hsp70 proteins are also involved in the response against oxidative stress (Azad et al., 2011;Shukla et al., 2014;Donovan and Marr, 2016) and heat-shock stress (Gong and Golic, 2006;Bettencourt et al., 2008;Shilova et al., 2018) in *Drosophila*, we postulated that increased Hsp70 production by decreased e-Ado signaling may also enhance the resistance against both stresses. To test this, we treated flies with either paraquat (a potent oxidative stress inducer; Fig. 4E) or a higher temperature (to induce heat-shock; Fig. 4F). We then compared the survival rate between the mutant flies and w^1118^ or Canton-S control flies. The results showed that *adoR* mutant flies were more resistant to paraquat and heat-shock treatment. Our results therefore demonstrate that the *Drosophila* AdoR pathway with its downstream gene *mod(mdg4)* suppresses Hsp70 protein production under a non-stress condition. Thus, the knockdown of *ent2*, *AdoR*, and *mod*(*mdg4*) results in increased levels of Hsp70, which in turn helps flies to respond to various stresses, including mHTT cytotoxicity, oxidative and heat-shock stresses (Fig. 4G).

## Discussion

Ado signaling represents an evolutionarily conserved pathway affecting a diverse array of stress responses (Fredholm, 2007). As a ubiquitous metabolite, Ado has evolved to become a conservative signal among eukaryotes. In previous studies, *Drosophila adoR* mutants (Dolezelova et al., 2007;Wu et al., 2009) and mice with a knockout of all four *adoRs* (Xiao et al., 2019) both displayed minor physiological alteration under normal conditions. This is consistent with the idea that Ado signaling more likely regulates the response to environmental changes (stresses) rather than being involved in maintaining fundamental homeostasis in both insect and mammalian models. Our study examined the impact of altering the expression of genes involved in Ado signaling and metabolism on the cytotoxicity and neurodegeneration phenotype of Q93 mHTT-expressing flies. We discovered a novel downstream target of this pathway, *mod(mdg4)*, and showed its effects on the downregulation of Hsp70 proteins, a well-known chaperone responsible for protecting cells against various stress conditions, including mHTT cytotoxicity, as well as thermal or oxidative stress (Soares et al., 2019).

The low level of Ado observed in our *da-Gal4* mHTT flies suggests that it might have a physiological role; lowering of the Ado level might represent a physiological response to cytotoxic stress. Consistently, our experimentally-decreased Ado signal rescued the mHTT phenotype, while an increased Ado signal had deleterious effects. Interestingly, a high level of Ado in the hemolymph has previously been observed in *Drosophila* infected by a parasitoid wasp (Novakova and Dolezal, 2011;Bajgar et al., 2015). A raised e-Ado titer has not only been shown to stimulate hemocyte proliferation in the lymph glands (Mondal et al., 2011), but also to trigger metabolic reprogramming and to switch the energy supply towards hemocytes (Bajgar et al., 2015). In contrast, our experiments show that a lowered e-Ado titer results in increased Hsp70 production. Increased Hsp70 has previously been shown to protect the cells from protein aggregates and cytotoxicity caused by mHTT expression, as well as some other challenges including oxidative stress (paraquat treatment) or heat-shock (Garbuz, 2017). The fine regulation of extracellular Ado in *Drosophila* might mediate the differential Ado responses via a single receptor isoform. Our earlier experiments on *Drosophila* cells also suggested that different cell types have different responses to Ado signaling (Fleischmannova et al., 2012).

Our data also showed that altered adenosine signaling through the receptor is closely connected to Ado transport, especially to ent2 transporter function. We observed that *adoR* and *ent2* knockdowns provide the most prominent rescue of mHTT phenotypes. In addition, the overexpression of *adoR* and *ent2* genes results in effects that are opposite to their knockdowns, thus supporting the importance of these genes as key regulators of mHTT phenotypes. Our previous report showed that responses to *adoR* and *ent2* mutations cause identical defects in associative learning and synaptic transmission (Knight et al., 2010). In the present study, we show that the phenotypic response of mHTT flies to *adoR* and *ent2* knockdowns are also identical. Our results suggest that the source of e-Ado for inducing AdoR signaling is mainly released by *ent2*. Consistently, the knockdown of *ent2* has previously been shown to block Ado release from *Drosophila* hemocytes upon an immune challenge (Bajgar et al., 2015), as well as from wounded cells stimulated by *scrib*-RNAi (Poernbacher and Vincent, 2018) or bleomycin feeding (Xu et al., 2020). These data support the idea that both *adoR* and *ent2* work in the same signaling pathway.

Our results revealed that lower AdoR signaling has a beneficial effect on mHTT-expressing flies, including increasing their tolerance to oxidative and heat-shock stresses. The effect of lower Ado signaling in mammals has been studied by pharmacologically blockading AdoRs, especially by the non-selective adenosine receptor antagonist caffeine. Interestingly, caffeine has beneficial effects on both neurodegenerative diseases and oxidative stress in humans (Rivera-Oliver and Diaz-Rios, 2014;Martini et al., 2016). In contrast, higher long-term Ado concentrations have cytotoxic effects by itself in both insect and mammalian cells (Schrier et al., 2001;Merighi et al., 2002). Chronic exposure to elevated Ado levels has a deleterious effect, causing tissue dysfunction, as has been observed in a mammalian system (Antonioli et al., 2019). Such disruption of nucleotide homeostasis has also been observed in mHTT-expressing R6/2 and Hdh150 mice (Toczek et al., 2016).

We identified a downstream target of the AdoR pathway, *mod(mdg4)*, which modulates mHTT cytotoxicity and aggregations. This gene has previously been implicated in the regulation of position effect variegation, chromatin structure, and neurodevelopment (Dorn and Krauss, 2003). The altered expression of *mod(mdg4)* has been observed in flies expressing untranslated RNA containing CAG and CUG repeats (Mutsuddi et al., 2004;van Eyk et al., 2011). In addition, *mod(mdg4*) has complex splicing, including *trans*-splicing, producing at least 31 isoforms (Krauss and Dorn, 2004). All isoforms contain a common N-terminal BTB/POZ domain which mediates the formation of homomeric, heteromeric, and oligomeric protein complexes (Bardwell and Treisman, 1994;Albagli et al., 1995;Espinas et al., 1999). Among these isoforms, only two [including *mod(mdg4)*-56.3 (isoform H) and *mod(mdg4)*-67.2 (isoform T)] have been functionally characterized. *mod(mdg4)*-56.3 is required during meiosis for maintaining chromosome pairing and segregation in males (Thomas et al., 2005;Soltani-Bejnood et al., 2007). *mod(mdg4)*-67.2 interacts with suppressor of hairy wing [Su(Hw)] and Centrosomal protein 190 kD (CP190) forming a chromatin insulator complex which inhibits the action of the enhancer on the promoter, and is important for early embryo development and oogenesis (Buchner et al., 2000;Soshnev et al., 2013;Melnikova et al., 2018). In the present study, we showed that *mod(mdg4)* is controlled by AdoR which consecutively works as a suppressor of Hsp70 chaperone. The downregulation of *adoR* or *mod(mdg4)* leads to the induction of Hsp70, which in turn suppresses mHTT aggregate formation and other stress phenotypes. Although our results showed that silencing all *mod(mdg4)* isoforms decreases cytotoxicity and mHTT inclusion formation, we could not clarify which of the specific isoforms is involved in such effects, since AdoR seems to regulate the transcriptions of multiple isoforms (Fig. S7). Further study will be needed to identify the specific *mod(mdg4)* isoform(s) connected to Hsp70 production.

In summary, our data suggest that the cascade (*ent2*)-*AdoR*-*mod(mdg4)*-*Hsp70* might represent an important general Ado signaling pathway involved in the response to various stress conditions, including reaction to mHTT cytotoxicity, oxidative damage, or thermal stress in *Drosophila* cells. The present study provides important insights into the molecular mechanisms of how Ado regulates mHTT aggregate formation and stress responses in *Drosophila*; this might be broadly applicable for understanding how the action of Ado affects disease pathogenesis.

## Materials and methods

### Fly stocks

Flies were reared at 25 °C on standard cornmeal medium. The following RNAi lines were acquired from the TRiP collection (Transgenic RNAi project) at Harvard Medical School: adgfA-Ri (BL67233), adgfC-Ri (BL42915), adgfD-Ri (BL56980), adenoK-Ri (BL64491), ent1-Ri (BL51055), adoR-Ri (BL27536), gfp-Ri (BL41552), mod(mdg4)-Ri (BL32995), cindr-Ri (BL38976), and ptp99A-Ri (BL57299). The following RNAi lines were acquired from the Vienna Drosophila RNAi Center (VDRC): ent2-Ri (ID100464), ent3-Ri (ID47536), cnt2-Ri (ID37161), and cg6184-Ri (ID107150).

Flies overexpressing human normal huntingtin (HTT) exon 1, Q20Httexon^1111F1L^, mutant pathogenic fragments (mHTT), Q93Httexon^14F132^ and elav^C155^-GAL4 were obtained from Prof. Lawrence Marsh (UC Irvine, USA) (Steffan et al., 2001). The UAS-overexpression lines, Ox-adenoK and Ox-adoR, were obtained from Dr. Ingrid Poernbacher (The Francis Crick Institute, UK) (Poernbacher and Vincent, 2018). gmr-GAL4 was obtained from Dr. Marek Jindra (Biology Centre CAS, Czechia). da-GAL4 was obtained from Dr. Ulrich Theopold (Stockholm University). The UAS overexpression strains Ox-adgfA, Ox-ent2, adoR^1^ and ent2^3^ mutant flies, were generated in our previous studies (Dolezal et al., 2003;Dolezal et al., 2005;Dolezelova et al., 2007;Knight et al., 2010).

### Eclosion rate and adult lifespan assay

For assessing the eclosion rate, male flies containing the desired RNAi or overexpression transgene (RiOx) in the second chromosome with genotype w^1118^/Y; RiOx /CyO; UAS-Q93/MKRS were crossed with females of *elav-GAL4*; +/+; +/+. The ratio of eclosed adults between *elav-GAL4*/+; RiOx/+; UAS-Q93/+ and *elav-GAL4*/+; RiOx/+; +/MKRS was then calculated. If the desired RiOx transgene was in the third chromosome, female flies containing *elav-GAL4*; +/+; RiOx were crossed with male w^1118^/Y; +/+; UAS-Q93/MKRS, and the ratio of eclosed adults between *elav-GAL4*; +/+; RiOx/UAS-Q93 and *elav-GAL4*; +/+; RiOx/MKRS was calculated.

For the adult survival assay, up to 30 newly-emerged female adults were placed in each cornmeal-containing vial and maintained at 25 °C. At least 200 flies of each genotype were tested and the number of dead flies was counted every day. Flies co-expressing RiOx and HTT Q20 were used for evaluating the effect of RNAi or overexpression of the desired transgenes.

### Extracellular adenosine and ATP level measurements

To collect the hemolymph, 6 third-instar larvae (96 hours post-oviposition) were torn in 150 μl of 1× PBS containing thiourea (0.1 mg/ml) to prevent melanization. The samples were then centrifuged at 5000× *g* for 5 min to separate the hemocytes and the supernatant was collected for measuring the extracellular adenosine or ATP level. For measuring the adenosine titer, 10 μl of hemolymph was mixed with the reagents of an adenosine assay kit (Biovision) following the manufacturer’s instructions. The fluorescent intensity was then quantified (Ex/Em = 533/587 nm) using a microplate reader (BioTek Synergy 4). For measuring the ATP level, 10 μl of hemolymph was incubated with 50 μl of CellTiter-Glo reagent (Promega) for 10 min. Then, the luminescent intensity was quantified using an Orion II microplate luminometer (Berthold). To calibrate the standard curve of ATP concentration, 25 μM ATP standard solution (Epicentre) was used for preparing a concentration gradient (0, 2, 4, 6, 8, 10 μM) of ATP solution and the luminescent intensity was measured for each concentration. The protein concentration of the hemolymph sample was determined by a Bradford assay. The adenosine and ATP concentrations were first normalized to protein concentration. Then, the values of Q20 and Q93 samples were normalized to values of the *GAL4* control sample. Six independent replicates for each genotype were performed for the analysis of adenosine and ATP levels.

### RNA extraction

The brains of 10 third-instar larvae (96 hours post-oviposition) or 15 whole female flies were pooled for each replicate. The samples were first homogenized in RiboZol (VWR) and the RNA phase was separated by chloroform. For brain samples, the RNA was precipitated by isopropanol, washed in 75% ethanol, and dissolved in nuclease-free water. For whole fly samples, the RNA phase was purified using NucleoSpin RNA columns (Macherey-Nagel) following the manufacturer’s instructions. All purified RNA samples were treated with DNase to prevent genomic DNA contamination. cDNA was synthesized from 2 μg of total RNA using a RevertAid H Minus First Strand cDNA Synthesis Kit (Thermo Fisher Scientific).

### Adenosine injection

Three- to five-day-old female adults were injected with 50 nl of 10 mM adenosine solution using a NANOJECT II (Drummond Scientific); control flies were injected with 50 nl of 1× PBS. Two hours post-injection, 15 injected flies for each replicate were collected for RNA extraction.

### Microarray analysis

The Affymetrix GeneChip® *Drosophila* genome 2.0 array system was used for microarray analysis following the standard protocol: 100 ng of RNA was amplified with a GeneChip 3′ express kit (Affymetrix), and 10 μg of labeled cRNA was hybridized to the chip according to the manufacturer’s instructions. The statistical analysis of array data was as described in our previous studies (Arefin et al., 2014;Kucerova et al., 2016). Storey’s q value (false discovery rate, FDR) was used to select significantly differentially transcribed genes (q < 0.05). Transcription raw data are shown in Table S2 and have been deposited in the ArrayExpress database (www.ebi.ac.uk/arrayexpress; accession No. E-MTAB-8699 and E-MTAB-8704).

### qPCR and primers

5× HOT FIREPol® EvaGreen® qPCR Mix Plus with ROX (Solis Biodyne) and an Eco Real-Time PCR System (Illumina) were used for qPCR. Each reaction contained 4 μl of EvaGreen qPCR mix, 0.5 μl each of forward and reverse primers (10 μM), 5 μl of diluted cDNA, and ddH2O to adjust the total volume to 20 μl. The list of primers is shown in Table S3. The expression level was calculated using the 2^-∆∆Ct^ method with the ct values of target genes normalized to a reference gene, ribosomal protein 49 (*rp49*).

### Imaging of retinal pigment cell degeneration

Twenty- and thirty-day-old female adults were collected and their eye depigmentation phenotypes were recorded. At least 30 individuals for each genotype were examined under a microscope, and at least five representative individuals were chosen for imaging. Pictures were taken with an EOS 550D camera (Canon) mounted on a SteREO Discovery V8 microscope (Zeiss).

### Brain immunostaining

Brains dissected from 10- or 20-day-old adult females were used for immunostaining. The brains were fixed in 4% PFA, permeabilized with PBST (0.1% Triton X-100), blocked in PAT (PBS, 0.1% Triton X-100, 1% BSA) and stained with antibodies in PBT (PBS, 0.3% Triton X-100, 0.1% BSA). Primary antibodies used in this study were mouse anti-HTT; MW8, which specifically binds to mHTT aggregates (1:40, DSHB); and rat anti-Elav (1:40, DSHB), which is a pan-neuronal antibody. Secondary antibodies were Alexa Fluor 488 anti-mouse and Alexa Fluor 647 anti-rat (1:200, Invitrogen). The samples were mounted in Fluoromount-G (Thermo Fisher Scientific) overnight, prior to image examination.

### Quantification of mHTT aggregates

Images of aggregates were taken using a FluoView 100 confocal microscope (Olympus). The intensity of mHTT aggregates detected by anti-HTT antibody (MW8) or anti-Elav were quantified using ImageJ software. The level of mHTT aggregates was quantified by normalizing the covering area of mHTT aggregate signal to Elav signal. At least six brain images from each genotype were analyzed.

### Western blot

Twenty heads, collected from 10-day-old adult females, were pooled for each replicate. The samples were homogenized in 100 μl of RIPA buffer with 1 μl of Halt™ proteinase inhibitor cocktail (Thermo Fisher Scientific). From each sample, 80 μl of supernatant was collected after 10 min of centrifugation at 12000× *g*, which was then mixed with 16 μl of 6× loading buffer. After boiling at 95 °C for 3 min, 10 μl were then loaded for running an SDS-PAGE gel. Proteins were then transfered to an Immobilon-E PVDF membrane (Millipore), which was then washed with 1× PBS containing 0.05% Tween 20 (three washes, each 15 min) and blocked in 5% BSA for 1 hour at room temperature before staining. The membrane was subsequently stained with primary antibodies overnight at 4 °C and secondary antibody for 1 hour at room temperature. After immunostaining, the membrane was treated with 2 ml of SuperSignal™ West Pico PLUS Chemiluminescent Substrate (Thermo Fisher Scientific) for 10 min at room temperature, and images were recorded using a Fujifilm LAS-3000 Imager. The primary antibodies used for staining were rat anti-Hsp70(7FB) (1:2000, Thermo Fisher Scientific) and mouse anti-Tub (1:500, DSHB). The secondary antibodies were donkey anti-rat IgG (H+L) HRP (1:5000, Thermo Fisher Scientific) and donkey anti-Mouse IgG (H+L) HRP (1:5000, Thermo Fisher Scientific).

### Paraquat injection

Three- to five-day-old males were collected for paraquat injection. Each fly was injected with 50 nl of 3 mM paraquat ringer solution using a NANOJECT II (Drummond Scientific). Control flies were injected with ringer buffer. Seventeen to twenty of injected flies were pooled into one vial for each replicate, and six replicates were performed for each treatment.

### Heat-shock treatment

The heat-shock procedure followed a previous study (Gong and Golic, 2006) with few modifications. Newly emerged males (0 or 1 day old) were collected and maintained on a standard cornmeal diet. The following day, ten flies were transferred into each empty vial and given a mild heat-shock at 35 °C for 30 min, then transferred to a circulating water bath at 39 °C. The number of surviving flies was checked every 10 min; flies which did not move any part of their body were considered dead.

### Statistical analysis

A Shapiro-Wilk test was applied to determine data normality. For data which were not normally distributed (*P* < 0.05), statistical significance was analyzed using the Mann-Whitney U-test. For normally distributed data (*P* > 0.05), statistical significance was established using Student’s t-test or one-way ANOVA with Tukey’s HSD *post-hoc* test. For the statistical analysis of survival curves, we used OASIS 2 to perform a weighted log-rank test (Han et al., 2016).

## Supporting information

Supplemental results

## Acknowledgements

We thank Dr. Ingrid Poernbacher (The Francis Crick Institute, UK), Prof. Lawrence Marsh (UC Irvine, USA), Dr. Marek Jindra (Biology Centre CAS, Czechia), Dr. Tomas Dolezal (University of South Bohemia, Czechia), Dr. Ulrich Theopold (Stockholm University), Bloomington *Drosophila* Stock Center and Vienna *Drosophila* Resource Center for providing us with fly strains. This work was supported by the grant agency of the University of South Bohemia (065/2017/P to Y-HL), junior grant project GACR (19-13784Y to LK) and European Community’s Programme Interreg Östereich-Tschechische Republik (REGGEN/ATCZ207 to MZ).

## Author Contributions

Y-HL performed the experiments and prepared the manuscript. HM assisted in recording the adult lifespan and eye phenotypes, and also as performed the brain dissection, immunochemistry and confocal microscopy imaging. LK performed the microarray sample preparation, analyzed the microarray data and paraquat injection. LR assisted in recording the adult lifespan and eye phenotypes, prepared fly strains and performed heat-shock treatment. TF established the methodologies for recording the eclosion rate and survival, and prepared fly strains. MZ conceived the project and supervised manuscript preparation.

