## Supplemental results for "Adenosine receptor and its downstream targets, mod(mdg4) and Hsp70, work as a signaling pathway modulating cytotoxic damage in *Drosophila*"

**Supplemental Information**

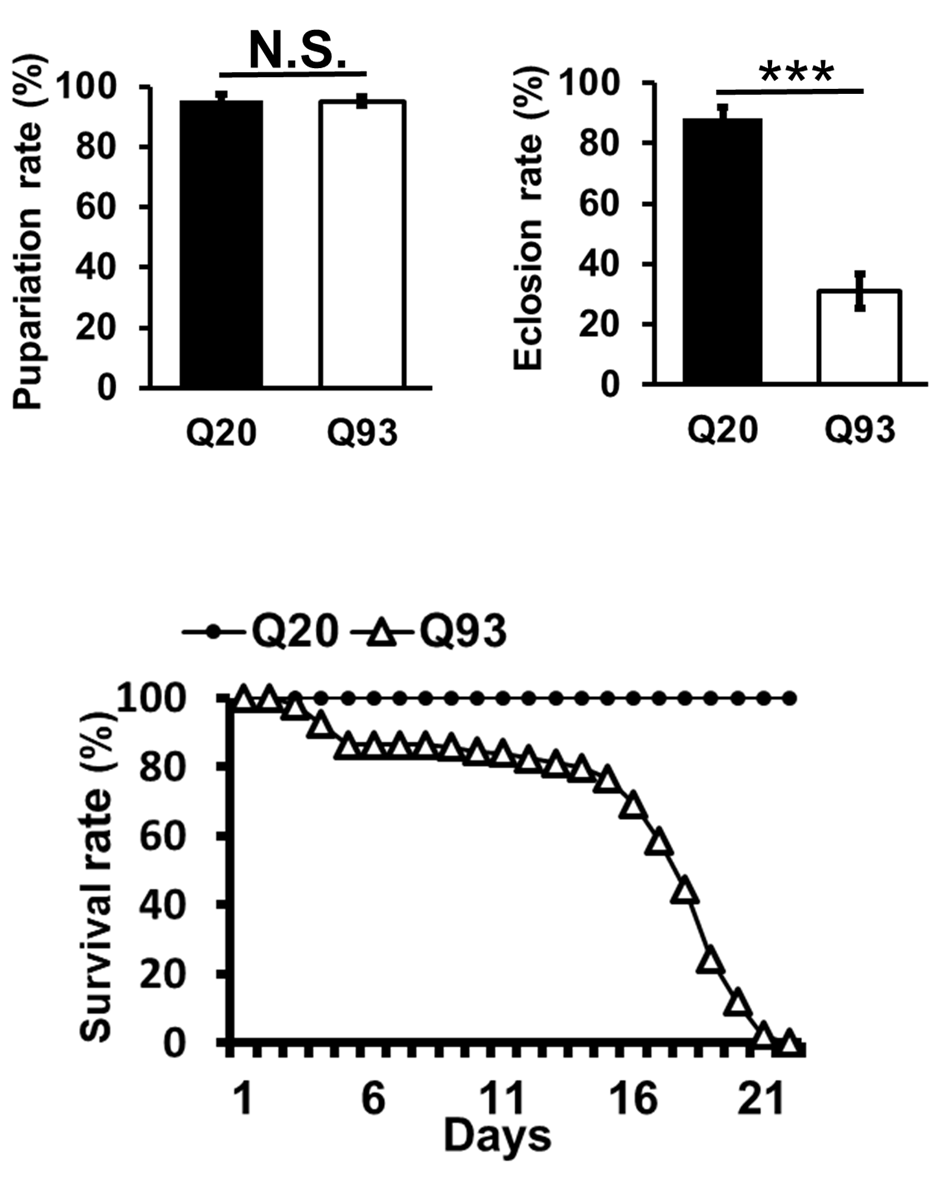

**Figure S1.** **mHTT expression decreased viability in *Drosophila*.** Flies expressing normal HTT (Q20) and mHTT (Q93) were driven by the pan-neuron driver (elav-GAL4), and the larval pupariation rate (A), adult eclosion rate (B), and adult survival (C) were recorded. At least five independent replicates were measured. Significance was examined using Student’s t-test: ****P* < 0.001; N.S., not significant. Error bars are presented as mean ± SEM

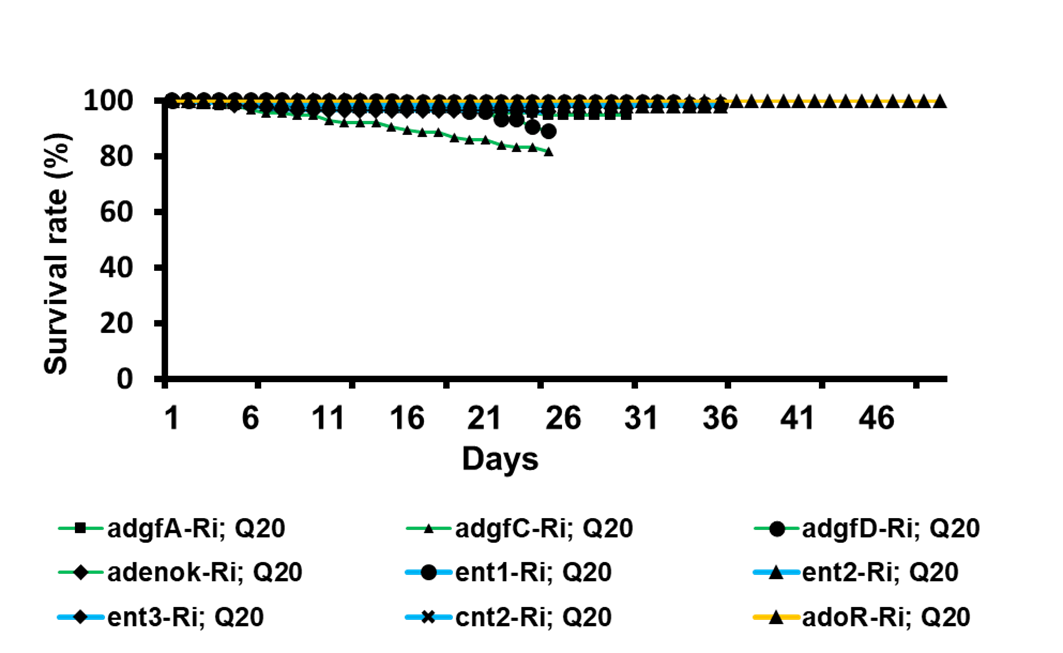

**Figure S2.** **RNAi of Ado metabolic, transport and receptor genes did not influence viability of flies.** Co-expression of normal Q20 HTT with RNAi of *adgf-A*, *adgf-C*, *adgf-D*, *adenoK*, *ent1*, *ent2*, *ent3*, *cnt2*, and *adoR* driven by the pan-neuronal driver (elav-GAL4). The number of dead flies was recorded until all corresponding experimental flies (expressing Q93 together with RNAi constructs) had died

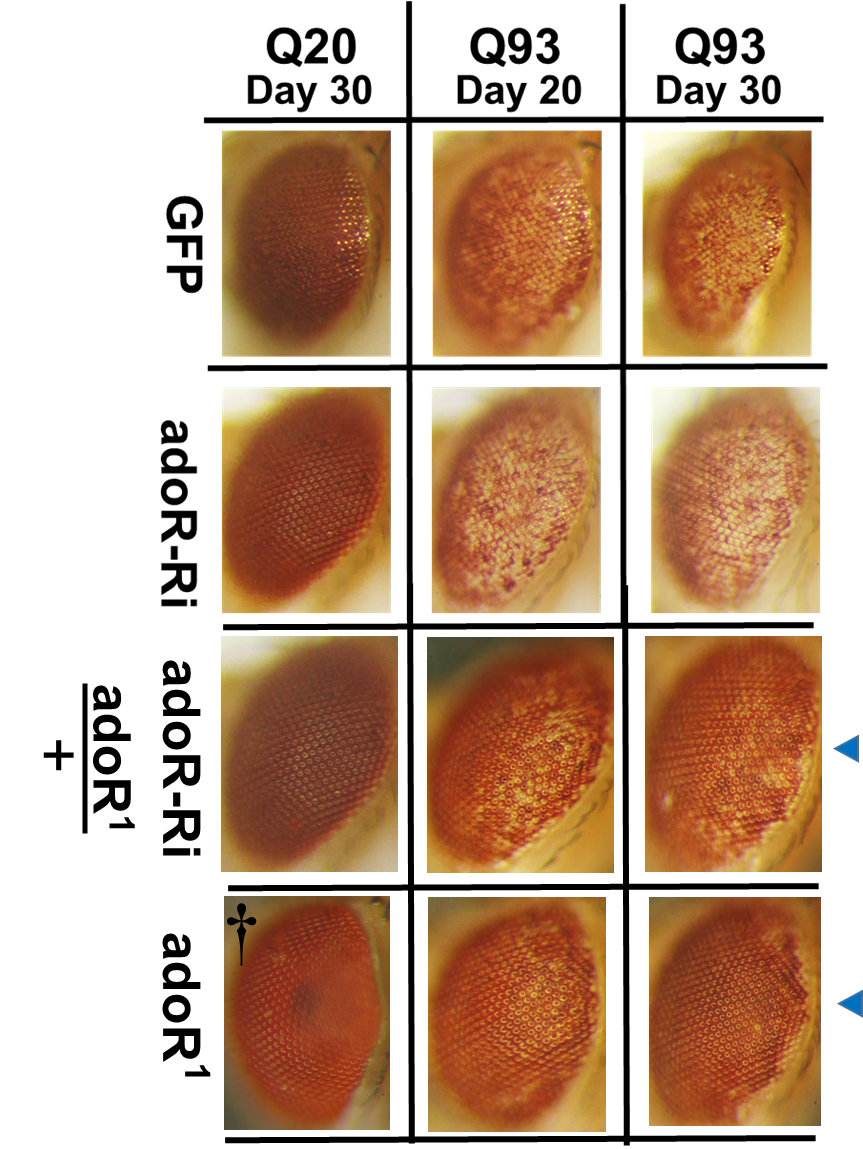

**Figure S3. Co-expression of Q93 with heterozygous adoR RNAi construct could not significantly rescue retinal pigment cell degeneration.** Retinal pigment cell degeneration in mHTT-expressing adult females (gmr>Q93) with RNAi silencing *adoR* or with *adoR* heterozygous and homozygous mutation

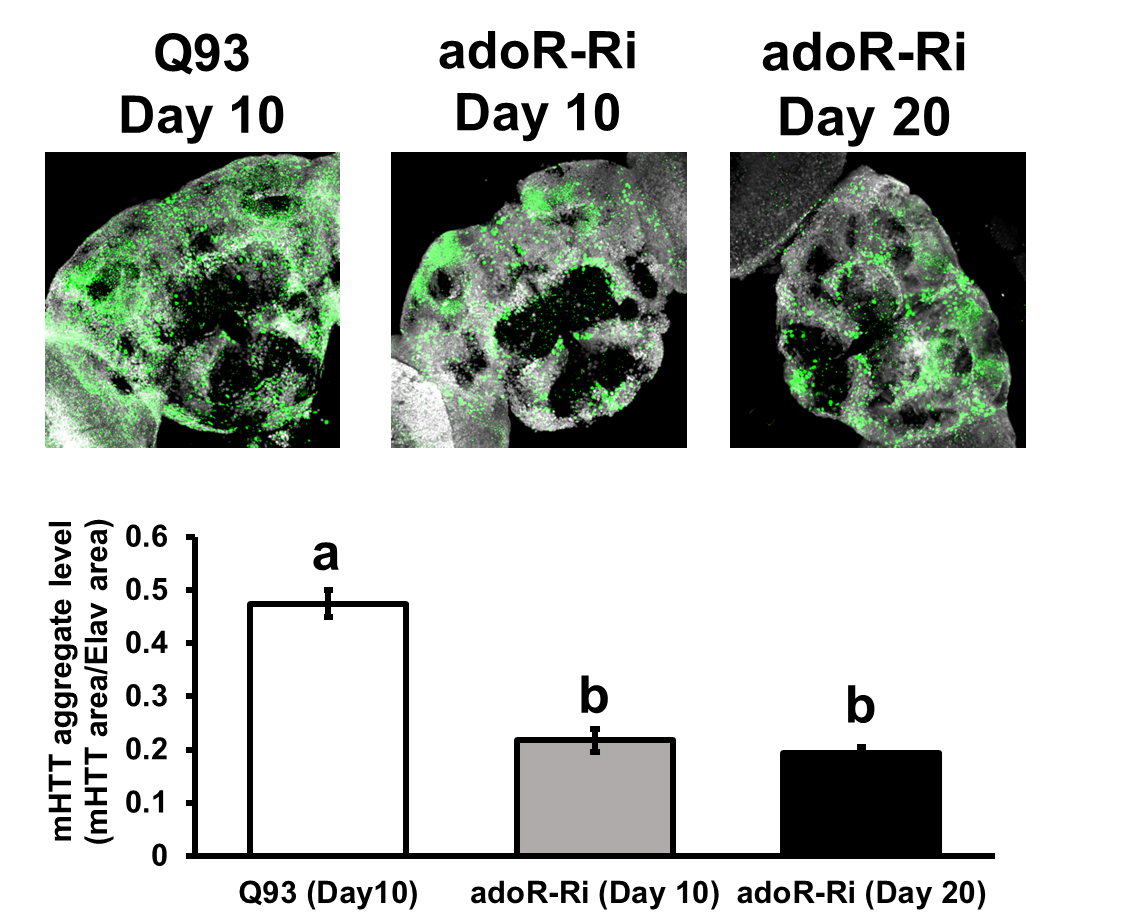

**Figure S4. AdoR RNAi suppressed mHTT aggregate formation in the brains of 10- and 20-day-old mHTT flies.** Representative confocal images of the brains of mHTT-expressing adult females (elav>Q93) with RNAi silencing *adoR*. Neuronal cells were detected with anti-Elav; mHTT aggregates were detected with anti-HTT (MW8). The level of mHTT aggregate formation was calculated by normalizing the area of mHTT signal to the area of Elav signal. Significance values of mHTT aggregates levels were analyzed by ANOVA; significant differences (*P* < 0.05) among treatment groups are marked with different letters. Error bars are presented as mean ± SEM. n = 6

| **probe ID** | **Gene name** | **Symbol** | **CG** | **Ent2^3^-L** | **AdoR^1^-L** | **AdoR^1^-A** | **Localization** |
| --- | --- | --- | --- | --- | --- | --- | --- |
| **1641190_at** | Jonah 65Aii | Jon65Aii | CG6580 | 2.79 | 2.5 | 3.18 | MG |
| **1627771_at** | --- |  | CG13075 | 2.58 | 2.91 | 1.93 | HG, MT |
| **1630590_at** | --- |  | CG18417 | 2.02 | 3.48 | 4.35 | MG, HG |
| **1624393_at** | white | w | CG2759 | 1.77 | 4.97 | 4.5 | MT |
| **1632003_a_at** | Protein tyrosine phosphatase 99A | Ptp99A | CG11516 | 1.53 | 1.41 | 2.32 | ID, NS, T |
| **1632404_at** | tetracycline resistance | rtet | CG5760 | 1.02 | 0.85 | 2.09 | MG, HG, I |
| **1633369_s_at** | --- |  | CG6184 | -0.23 | -0.21 | -0.68 | NS, tes |
| **1637389_at** | Ionotropic receptor 100a | Ir100a | CG11575 | -0.67 | -0.68 | -2.61 | non-spec |
| **1635901_at** | [Protein S-acyltransferase](http://www.genecards.org/cgi-bin/carddisp.pl?gene=CG17197) |  | CG17197 | -0.85 | -0.63 | -1.09 | ID, tes |
| **1624020_at** | [modifier of mdg4](http://www.genecards.org/cgi-bin/carddisp.pl?gene=mod(mdg4)) | mod(mdg4) | CG32491 | -1 | -1.4 | -0.57 | NS, ID |
| **1640850_at** | CIN85 and CD2AP orthologue | cindr | CG31012 | -1.11 | -1.1 | -1.02 | NS, SS, ID, tes |
| **1627953_at** | modifier of mdg4 | mod(mdg4) | CG32491 | -1.28 | -1.08 | -1.19 | NS, ID |
| **1633748_at** | NADH dehydrogenase (ubiquinone) 20 kDa subunit-like | ND-20L | CG2014 | -1.94 | -2.05 | -0.82 | ID, tes |

**Table S1. Intersection genes between the three datasets of microarray analysis.** Genes highlighted in red and blue were significantly up- and downregulated (q<0.05), respectively, in the larvae of *ent2* (Ent2^3^-L) mutant and *adoR* mutants (AdoR^1^-L) as well as adult of *adoR* mutant (AdoR^1^-A). Tissue localization of each gene expression was obtained from Flybase (http://flybase.org/). Tissue abbreviations: midgut (MG), hindgut (HG), Malpighian tubule (MT), imaginal disc (ID), integument (I), sensory system (SS), nervous system (NS), trachea (T), testis (tes), non-specific expression (non-spec)

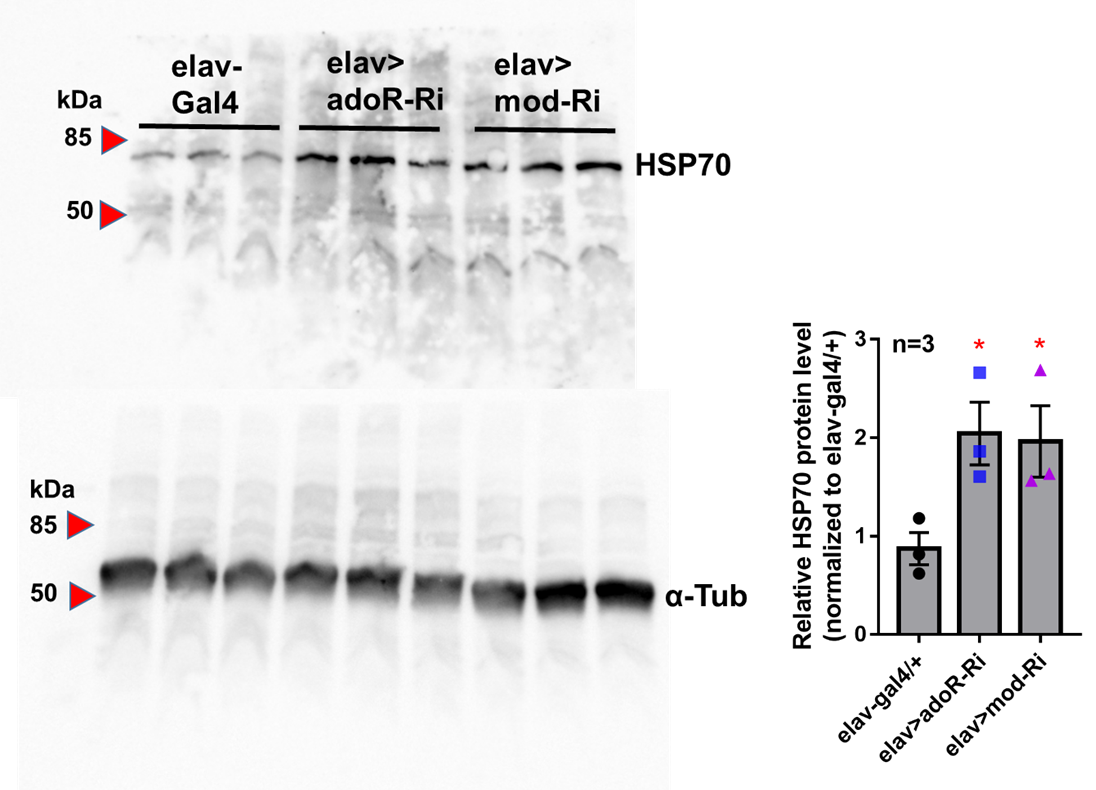

**Figure S5. Full image of western blot results shown in Figure 4A.** Hsp70 protein levels in the heads of 10-day-old adult females with RNAi silencing adoR (elav>adoR-Ri), mod(mdg4) (elav>mod-Ri) and control (elav-gal4/+). The Hsp70 protein level was quantified by normalizing the intensity of the Hsp70 band to the α-Tubulin band using ImageJ; values of RNAi treatment groups were further normalized to the elav-gal4 control. Significance was analyzed by Student’s t-test; significant differences between the control and each RNAi treatment group are labeled as **P* < 0.05. Error bars are presented as mean ± SEM. n = 3

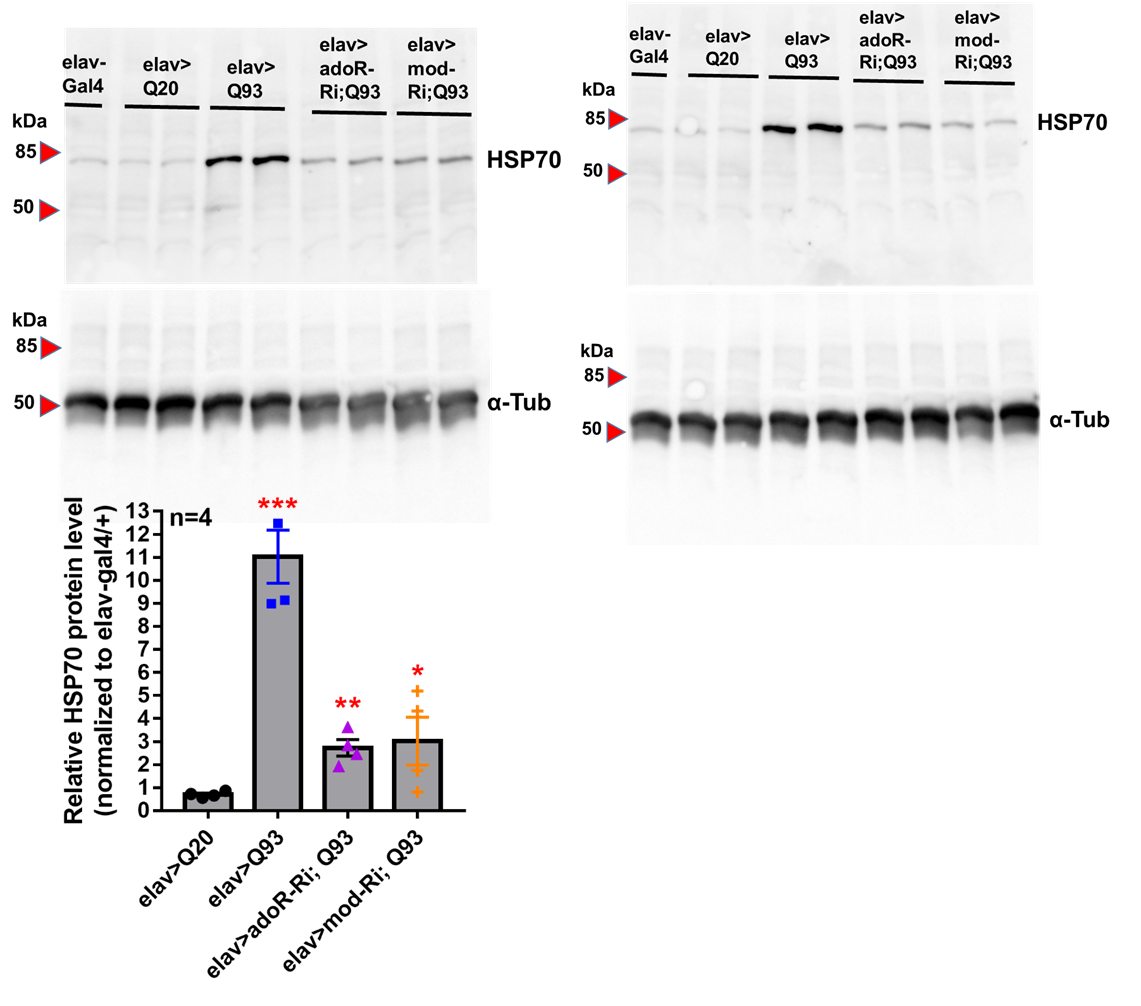

**Figure S6. Full image of western blot results shown in Figure 4C.** Hsp70 protein levels in the heads of 10-day-old HTT (elav>Q20) or mHTT-expressing (elav>Q93) adult females with RNAi silencing adoR and mod(mdg4). The Hsp70 protein level was quantified by normalizing the intensity of the Hsp70 band to the α-Tubulin band using ImageJ; values for each treatment group were further normalized to the elav-gal4 control. Significance was analyzed by Student’s t-test; significant differences between HTT-expressing flies (elav>Q20) with each RNAi treatment of Q93 expressing flies are labeled as follows: **P* < 0.05, ***P* < 0.01, ****P* < 0.001. Error bars are presented as mean ± SEM. n = 4

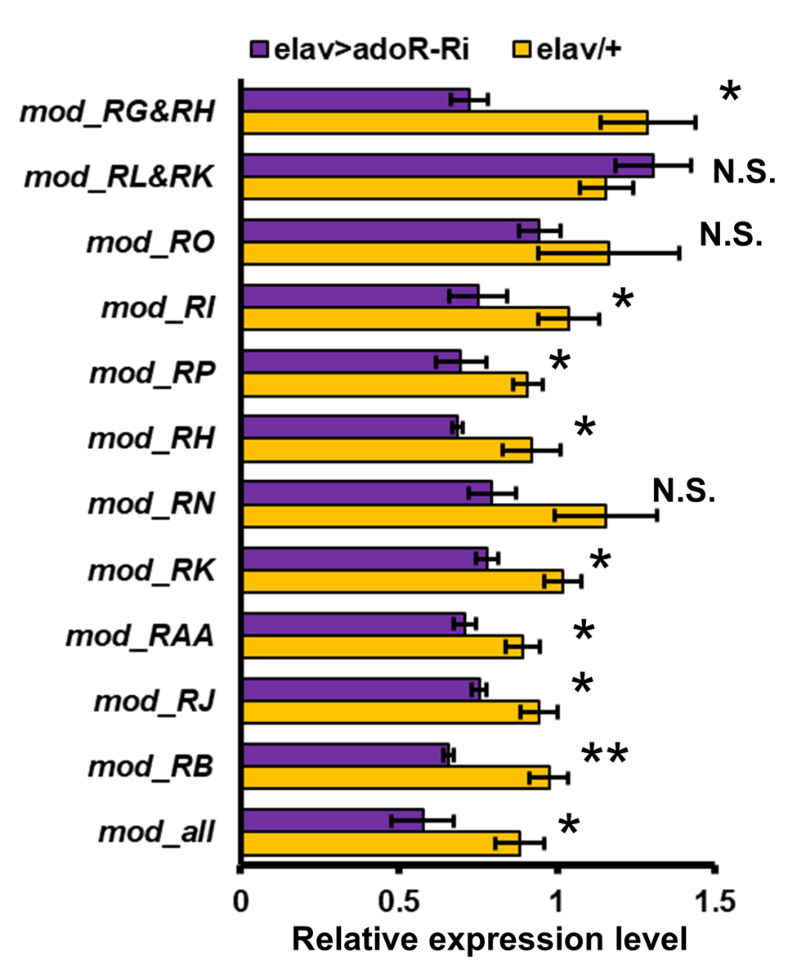

**Figure S7. Multiple isoforms of *mod(mdg4)* were downregulated by adoR RNAi.** *adoR* RNAi transgene (adoR-Ri) expression was driven by the pan-neuronal driver (elav>adoR-Ri); control flies contained only *elav*-GAL*4* (elav/+). Mod_all indicates that the primers targeted all *mod(mdg4)* isoforms. Isoforms L and G do not have their own unique exonal region, therefore it is possible for the qPCR primers to target two isoforms simultaneously (presented as RG&RG and RL&RK). qPCR result significance was examined using Student’s t-test: **P* < 0.05; ***P* < 0.01; N.S., not significant
